# Improving Designer Glycan Production in *Escherichia coli* through Model-Guided Metabolic Engineering

**DOI:** 10.1101/160853

**Authors:** Joseph A. Wayman, Cameron Glasscock, Thomas J. Mansell, Matthew P. DeLisa, Jeffrey D. Varner

## Abstract

Asparagine-linked (*N*-linked) glycosylation is the most common protein modification in eukaryotes, affecting over two-thirds of the proteome. Glycosylation is also critical to the pharmacokinetic activity and immunogenicity of many therapeutic proteins currently produced in complex eukaryotic hosts. The discovery of a protein glycosylation pathway in the pathogen *Campylobacter jejuni* and its subsequent transfer into laboratory strains of *Escherichia coli* has spurred great interest in glycoprotein production in prokaryotes. However, prokaryotic glycoprotein production has several drawbacks, including insufficient availability of non-native glycan precursors. To address this limitation, we used a constraint-based model of *E. coli* metabolism in combination with heuristic optimization to design gene knockout strains that overproduced glycan precursors. First, we incorporated reactions associated with *C. jejuni* glycan assembly into a genome-scale model of *E. coli* metabolism. We then identified gene knockout strains that coupled optimal growth to glycan synthesis. Simulations suggested that these growth-coupled glycan overproducing strains had metabolic imbalances that rerouted flux toward glycan precursor synthesis. We then validated the model-identified knockout strains experimentally by measuring glycan expression using a flow cytometric-based assay involving fluorescent labeling of cell surface-displayed glycans. Overall, this study demonstrates the promising role that metabolic modeling can play in optimizing the performance of a next-generation microbial glycosylation platform.

## 1. Introduction

Protein glycosylation is the attachment of glycans (mono-, oligo-, or polysaccharide) to specific amino acid residues in proteins, most commonly asparagine (*N*-linked) or serine and threonine (*O*-linked) residues. Roughly three-quarters of eukaryotic proteins and more than half of prokaryotic proteins are glycosylated [1]. Glycosylation is also vitally important to the development of many protein biologics, and has been harnessed for enhancing therapeutic properties such as half-life extension [2, 3, 4, 5], antibody-mediated cytotoxicity [6, 7], and immunogenicity [8, 9, 10].

Though once thought to occur only in eukaryotes, protein glycosylation has now been discovered in all three domains of life, including bacteria [11]. The best characterized bacterial *N*-glycosylation system is that of the human pathogen *Campylobacter jejuni* [12]. The *C. jejuni* glycan has the form of a branched heptasaccharide Glc GalNAc_5_ Bac, where Glc is glucose, GalNAc is *N*-acetylgalactosamine, and Bac is bacillosamine. This glycan is assembled on the lipid carrier undecaprenyl pyrophosphate (Und-PP) on the cytoplasmic face of the inner membrane by an enzymatic pathway encoded by the *pgl* (protein glycosylation) locus (Fig. 1). The fully assembled glycan is flipped across the membrane and transferred to asparagine residues in acceptor proteins by the oligosaccharyltransferase (OST) PglB. PglB attaches the heptasaccharide to periplasmically-localized proteins containing the consensus sequence D/E-X-N-Z-S/T, where X and Z are any residue except proline [13, 14].

**Figure 1:**
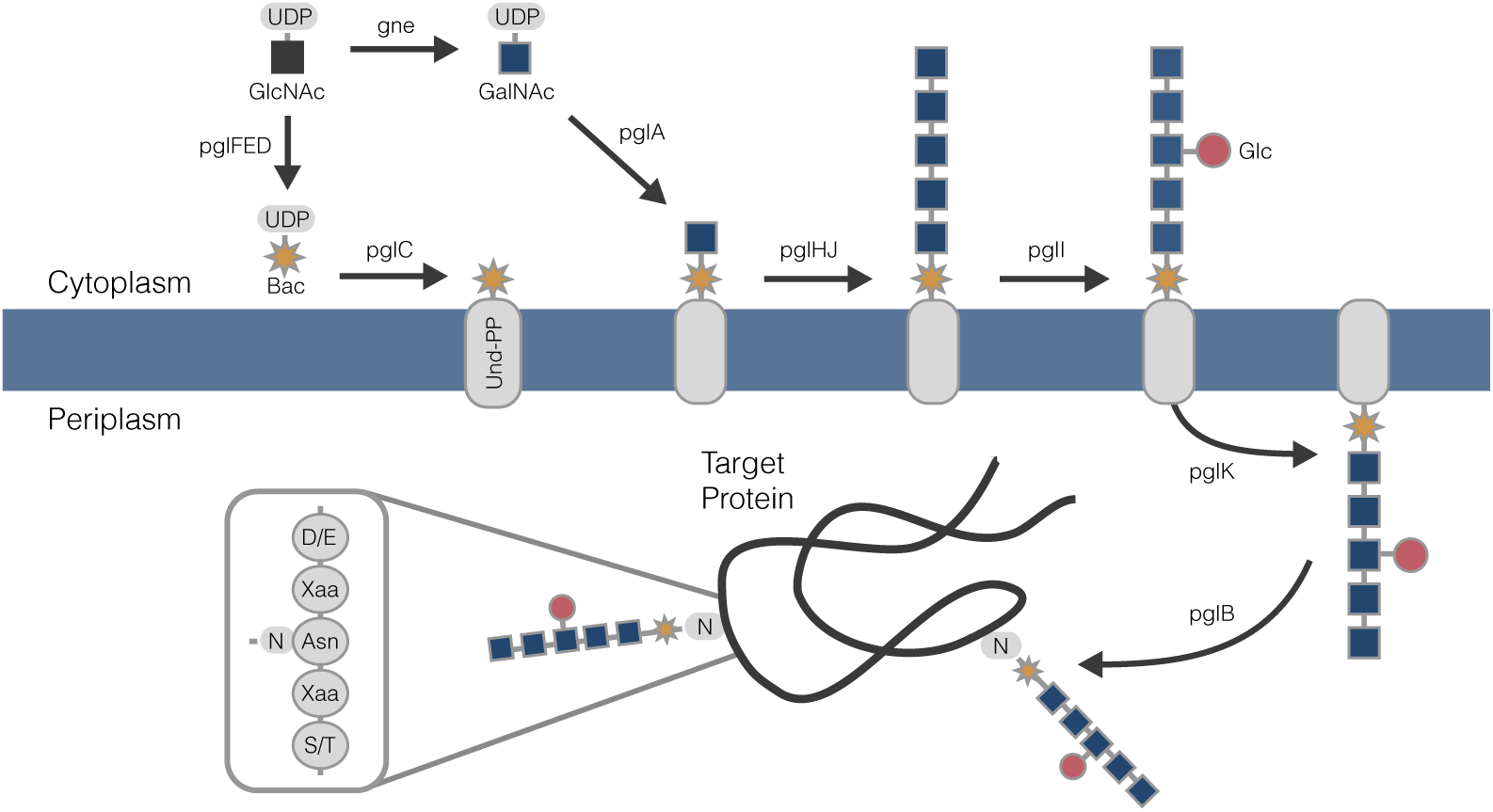
Glycosylation pathway in *C. jejuni* and *E. coli*. Glycan assembly, facilitated by *pgl* locus enzymes, takes place on a lipid carrier, undecaprenyl pyrophosphate (Und-PP), from cytoplasmic pools of nucleotide-activated sugars N-acetylglucosamine (GlcNAc), N-acetylgalactosamine (GalNAc), and glucose (Glc). The glycan is then flipped onto the periplasmic side of the inner membrane, where it is transferred to an asparagine residue on a glycoprotein acceptor motif.

The functional transfer of this system into *E. coli* [15] has spurred interest in recombinant production of glycans and ultimately therapeutic glycoproteins in this genetically tractable bacterial host [16, 17]. Along these lines, glycosylation-competent *E. coli* cells have been used to produce a variety of periplasmic and extracellular glycoproteins including antibodies [13] and conjugate vaccine candidates [18]. The promiscuity of the PglB enzyme towards structurally diverse lipid-linked glycan substrates has been exploited to further expand the *E. coli* platform, enabling the creation of glycoproteins bearing different bacterial O-polysaccharide antigens [18, 19] and even the eukaryotic trimannosyl core *N*-glycan produced by a synthetic pathway comprised of four yeast glycosyltransferases [20]. However, while PglB can efficiently glycosylate native *C. jejuni* acceptor proteins with cognate Glc GalNAc_5_ Bac glycan in engineered *E. coli*, glycosylation of non-Campylobacter target proteins is often much less efficient [21], especially in combination with heterologous glycan structures [20].

In engineered *E. coli*, protein glycosylation is affected by the availability of lipid carriers, and the availability of nucleotide-activated sugar substrates serving as glycan precursors [16, 22]. Hence, a plausible strategy for increasing glycosylation efficiency is to optimize the levels of these key reaction intermediates and their related biosynthetic pathways. Along these lines, Wright and coworkers applied genome-scale metabolic engineering techniques to improve glycosylation efficiency in *E. coli*. Using a high-throughput proteomic screening and probabilistic metabolic network analysis, they showed that upregulation of the glyoxylate cycle by overexpression of isocitrate lyase (*aceA* / *icl*) increased glycosylation efficiency of a prototypic protein by three-fold [23]. Further, genome-wide screening of gene overexpression identified targets that increased glycoprotein production as well as glycosylation efficiency [24]; genes in pathways associated with glycan precursor synthesis (UDP-GlcNAc) as well as lipid carrier production (isoprenoid synthesis) were identified as bottlenecks. Improved glycosylation efficiency has also been achieved by supplementing growth media with GlcNAc [25] or increasing the expression of PglB via codon optimization [26]. These studies and others have demonstrated the complex interplay between recombinant protein production, glycan synthesis and assembly, and glycosylation efficiency.

In this study, we addressed one of the challenges facing high-level glycoprotein production in engineered *E. coli*, namely the availability of glycan precursors, using constraint-based modeling. In particular, we used a constraint-based model of *E. coli* metabolism, in combination with heuristic optimization, to design gene knockout strains that overproduced glycan precursors. First, we incorporated reactions associated with *C. jejuni* glycan assembly into a genome-scale model of *E. coli* metabolism. We then used a combination of constraint-based modeling and simulated annealing to identify gene knockout strains that coupled optimal growth to glycan synthesis. Simulations suggested that these growth-coupled glycan overproducing strains had metabolic imbalances that rerouted flux toward glycan precursor synthesis. We then experimentally validated the model-identified metabolic designs using a flow cytometric-based assay for quantifying cellular *N*-glycans in *E. coli* [20]. Consistent with simulations, the best model-predicted changes increased glycan production by nearly 3-fold compared with the glycan production level in wild-type (wt) *E. coli* cells. Taken together, our results reveal the significant impact that metabolic modeling can have on designing chassis strains with enhanced *N*-linked protein glycosyation capabilities.

### 1.1 Results

#### Construction of a constraint-based model of N-linked glycosylation in E. coli

A constraint-based model of *N*-glycosylation in *E. coli* was used to identify genetic knockouts that coupled glycan biosynthesis with optimal growth. We augmented the existing genome-scale *E. coli* model iAF1260 from Palsson and coworkers [27] to include the reactions of the *C. jejuni* glycosylation pathway (Table 1). The adapted network consisted of 2395 reactions, 1271 open reading frames, and 1986 metabolites segregated into cytoplasmic, periplasmic, and extracellular compartments. Added reactions included the biochemical transformations catalyzed by the glycosyltransferases (e.g., PglA, PglC) associated with glycan biosynthesis, PglK flippase-mediated translocation of the glycan into the periplasm, and PglB-mediated glycan conjugation to an acceptor protein (Fig. 1). In addition, we incorporated the transcriptional regulatory network of Covert *et al.*, consisting of 101 transcription factors, regulating the state of the metabolic genes [28]. This regulatory network imparts Boolean constraints on metabolic fluxes based upon the nutrient environment. The model code is available for download under an MIT software license from the Varnerlab website [29].

**Table 1:**
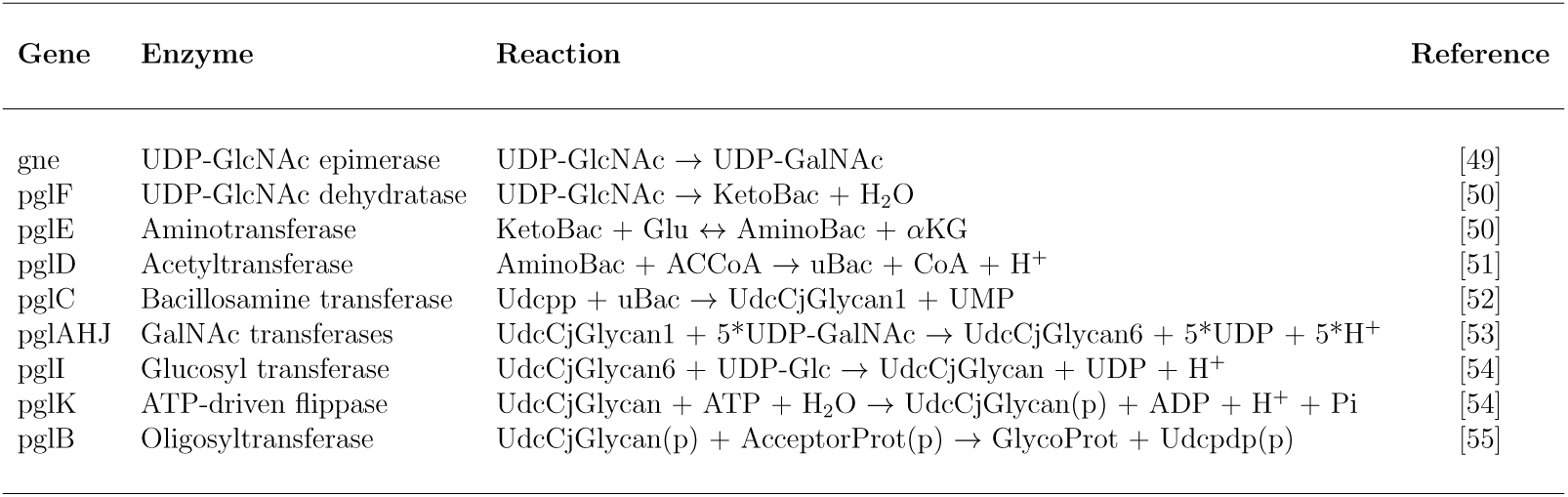
Reactions added to the *E. coli* model iAF1260 [27] for biosynthesis of *C. jejuni* glycan. Species localized to the periplasm are denoted by (p), all others are cytoplasmic. Abbreviations: UDP-N-Acetyl-D-Glucosamine, UDP-GlcNAc; UDP-N-Acetyl-D-Galactosamine, UDP-GalNAc; UDP-2-acetamido-2,6-dideoxy-*α*-D-xylo-4-hexulose, KetoBac; L-Glutamate, Glu; UDP-N-Acetylbacillosamine, AminoBac; *α*-ketoglutarate, *α*KG; Acetyl-CoA, ACCoA; UDP-N,N’-diacetylbacillosamine, uBac; Coenzyme A, CoA; Undecaprenyl phosphate, Udcpp; *C. jejuni* glycan intermediates, UdcCjGlycan1, UdcCjGlycan6; Uridine monophosphate, UMP; Uridine diphosphate, UDP; UDP-Glucose, UDP-Glc; Lipid-linked *C. jejuni* glycan, UdcCjGlycan; Acceptor protein, AcceptorProt; GlycoProt, Glycoprotein; Undecaprenyl diphosphate, Udcpdp.

#### Identification of growth-coupled gene knockout strains

To identify genetic knockouts that coupled optimal growth to glycan biosynthesis, we used heuristic optimization and the constraint-based model (see Materials and Methods). Coupling growth to glycan synthesis was desirable for several reasons. Foremost amongst these, growth-coupled strains create stoichiometric imbalances that reroute metabolic flux toward the desired product as a consequence of growth [30, 31]. Therefore, faster growth requires increased glycan formation. Thus, optimizing glycan production through adaptive evolution is made trivial by selecting for growth through serial passage [31, 32]. Several methods have been developed to estimate genetic knockouts using constraint-based models. In this study, we used simulated annealing to search over the states of metabolic enzyme and transcription factor (TF) genes to identify the desired phenotype (Fig. 2). The state of each gene was represented as a binary array, where a one indicated normal activity, while a zero indicated a genetic knockout or regulatory repression. Boolean rules informed by nutrient conditions controlled the TF genes, which in turn controlled the state of the metabolic genes. Once defined, the genetic state of the model modified the flux constraints placed on each reaction. For example, the reaction governed by pyruvate dehydrogenase, a multi-component enzyme, relied on the assembly of three enzymes: AceE, AceF, and Lpd. This reaction was encoded as:

**Figure 2:**
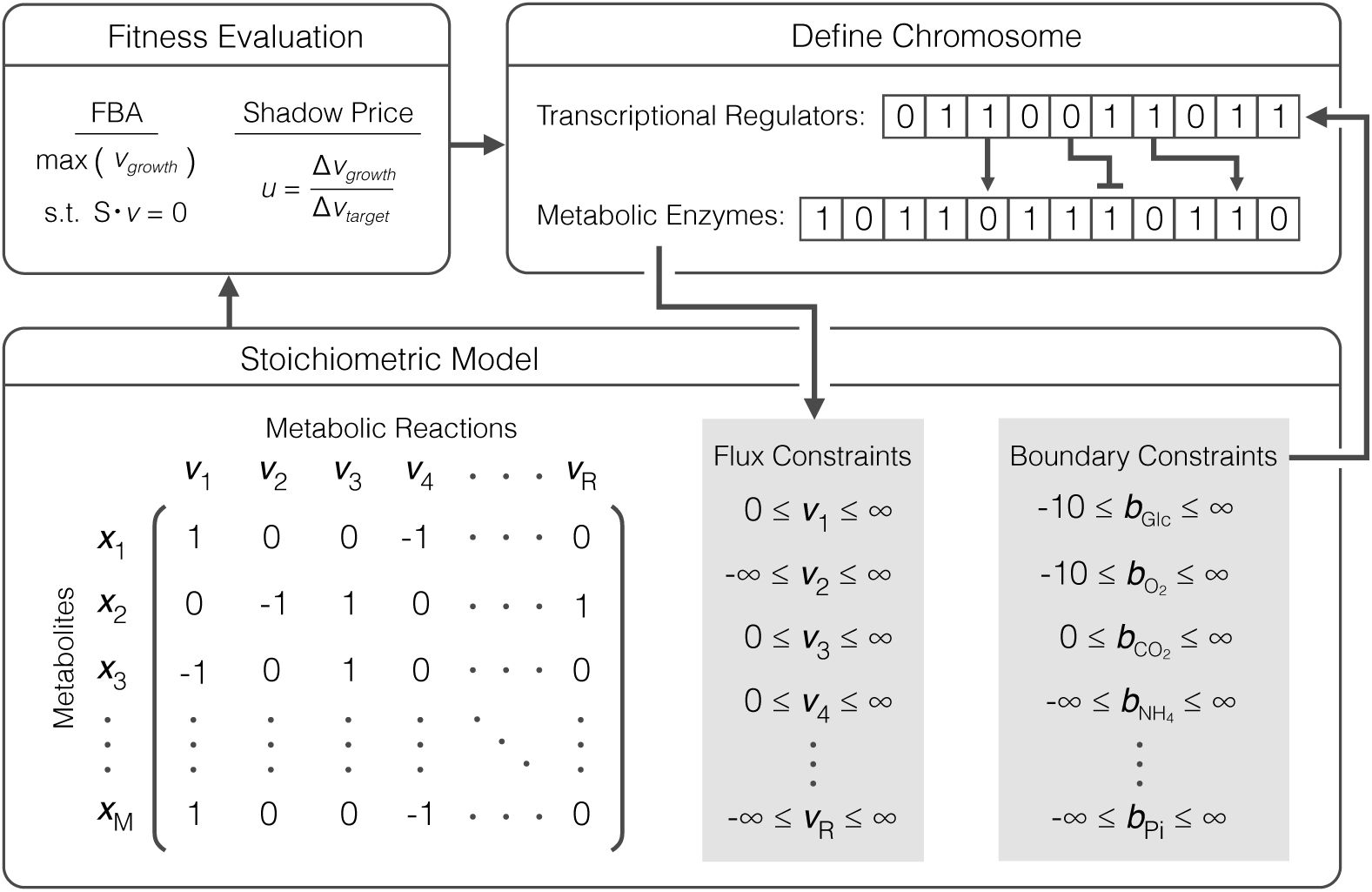
Heuristic optimization approach used to identify strains coupling growth to glycan production. The chromosome is defined as two separate binary arrays, one defining the state of metabolic enzyme expression and another defining the state of transcriptional regulator activation. Gene repression and knockouts are designated by zeros. Nutrient conditions define the boundary constraints within the stoichiometric model which in turn affect the state of the metabolic enzyme chromosome. Gene repression and knockouts determine the constraints placed on fluxes in the stoichiometric model. Nutrients are mapped to the state of transcriptional regulators and genes are mapped to the state of flux constraints using Boolean rules as defined in [27, 28]. Flux balance analysis is used to maximize growth rate under the constraints imposed by the mutant strain and transcriptional regulation and the fitness objective is calculated. Here, we use shadow price; the strain is accepted or rejected based on the change in fitness and a Boltzmann criterion. New mutant strains are randomly generated from accepted ones. The search continues until a positive shadow price is achieved.

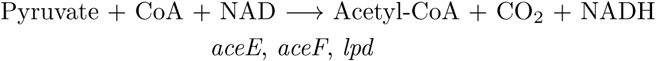

Thus, if any of the genes *aceE, aceF*, or *lpd* was knocked out or transcriptionally repressed, the flux through this reaction was bound to zero. Gene-protein-reaction (GPR) associations from the iAF1260 network were used in this study [27]. The simulated annealing algorithm performed a random search of genetic knockouts, iteratively applying flux constraints based on the genetic state, then performing a flux balance analysis simulation. To identify growth-coupled glycan producing strains, we optimized the shadow price given by:

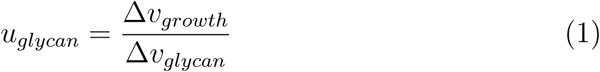

where Δ*v*_*growth*_ denotes the change in growth rate for a forced change in glycan flux Δ*v*_*glycan*_, and *v*_*glycan*_ denotes the flux representing the fully assembled *C. jejuni* glycan flipped into the periplasm. The shadow price *u*_*glycan*_ was calculated for a particular knockout strain by first calculating the optimal growth with the glycan flux constrained to zero. A second simulation was then performed with a forced incremental change in the glycan flux in order to obtain the difference in growth rate. The search algorithm continued until *u*_*glycan*_ *>* 0, indicating a growth-coupled phenotype (Supplementary Fig. S1).

We identified growth-coupled knockout strains with four or fewer knock-outs for growth on glucose as the sole source of carbon and energy (Table 2). We performed optimization simulations using boundary conditions representing minimal medium with a single 6-, 5-, and 3-carbon substrate. A well-defined minimal media allowed for precise control over nutrient conditions experimentally, and was accurately simulated, particularly for the transcriptional regulatory network. For each substrate, we performed ten independent optimization simulations to identify growth-coupled strains. We considered growth-coupled strains with four or fewer knockouts (those most likely to be experimentally viable) by restricting the formation of extracellular byproducts to acetate. For example, for *E. coli* glycosylation mutant 2 (EcGM2; *E. coli* iAF1260 Δ*sdh* Δ*gnd* Δ*pta* Δ*eutD*), the strain with the highest simulated glycan yield, the optimal growth rate occurs at a non-zero glycan flux (Fig. S1B). All growth-coupled strains contained a knockout of succinate dehydrogenase (*sdh*) and truncated pentose phosphate pathway (PPP) flux at either glucose 6-phosphate-1-dehydrogenase (*zwf*), 6-phosphogluconolactonase (*pgl*), or 6-phosphogluconate dehydrogenase (*gnd*).

**Table 2:**
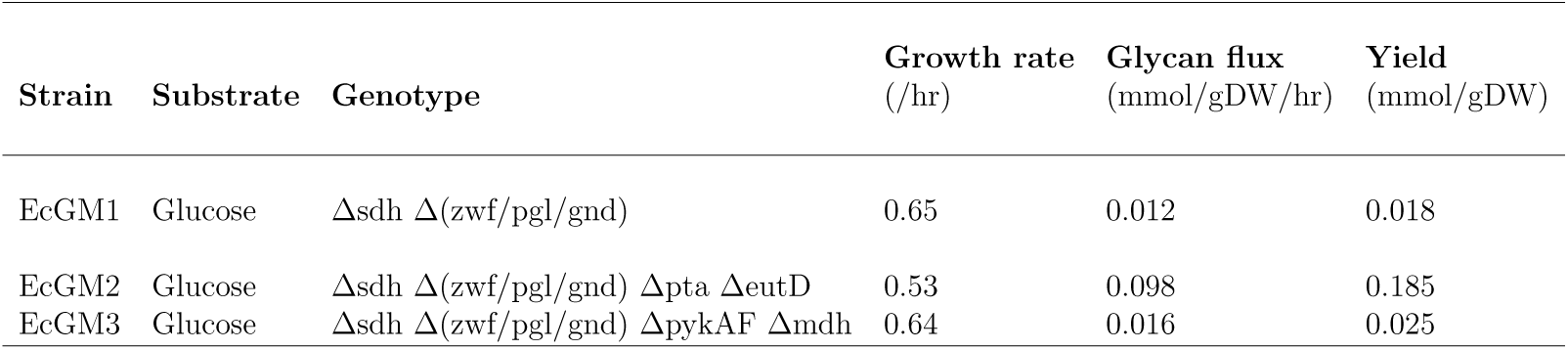
Growth-coupled strains producing *C. jejuni* glycan identified by flux balance analysis and heuristic optimization using single carbon substrate. Knockouts listing multiple genes indicate that knockout of any one of those genes produces the same phenotype in the model. Abbreviations: D-Glucose, Glc; *E. coli* glycosylating mutant, EcGM

| Strain | Substrate | Genotype | Growth rate<br>(/hr) | Glycan flux<br>(mmol/gDW/hr) | Yield<br>(mmol/gDW) |
| --- | --- | --- | --- | --- | --- |
| EcGM1 | Glucose | Δsdh $\Delta$(zwf/pgl/gnd) | 0.65 | 0.012 | 0.018 |
| EcGM2 | Glucose | Δsdh $\Delta$(zwf/pgl/gnd) Δpta ΔeutD | 0.53 | 0.098 | 0.185 |
| EcGM3 | Glucose | Δsdh $\Delta$(zwf/pgl/gnd) ΔpykAF Δmdh | 0.64 | 0.016 | 0.025 |

### Flux analysis of N-glycan production in growth-coupled strains

Growth-coupled glycan producing strains had increased glycolytic flux, and decreased amino acid biosynthesis compared to glycan production in the wt strain background (Fig. 3). We compared the normalized flux values for EcGM2 with the wt strain. Normalizing all fluxes to glucose uptake rate, EcGM2 displayed greater flux through glycolysis by cutting off the PPP via knockout of NADPH-producing *gnd* (Fig. 3A). EcGM2 also had decreased synthesis of every amino acid except for glutamine, indicating a source of stoichiometric imbalance that may be relieved by synthesis of the glycan precursor UDP-GlcNAc. Further, the PEP-pyruvate node acted as a switch point in central carbon metabolism (Fig. 3B). Here, PEP and pyruvate, the products of glycolysis, enter the TCA cycle through decarboxylation of pyruvate to acetyl-CoA (ACCoA) and carboxylation of PEP to form oxaloacetate (OAA) [33]. The latter replenishes TCA cycle intermediates that exited TCA for anabolic processes. EcGM2, with a diminished anabolic capacity for cell growth, displayed lower flux through PEP carboxylase (*ppc*). However, as the result of high glycolytic flux, EcGM2 had increased flux through pyruvate dehydrogenase (*aceEF*), sending carbon into the oxidative branch of the TCA cycle. It is known that high glucose uptake rates result in excess acetyl-CoA, surpassing the capacity of the TCA cycle. Because of this excess flux, wt *E. coli* grown on glucose commonly displays acetate fermentation, even under aerobic conditions [34]. We observed increased acetate secretion in EcGM1 simulations, but through a route differing from wild-type cells. The knockouts Δ*pta* and Δ*eutD* prevented ATP-generating acetate secretion. Flux was instead routed through the redox-neutral reactions initiated by acetaldehyde dehydrogenase (*mhpF*). Excess acetyl-CoA was also utilized in the pathway generating UDP-GlcNAc. Lastly, EcGM2 displayed a shift in cofactor production (Fig. 3C). Higher flux through glycolysis naturally led to NADH overproduction. On the other hand, the primary source of NADPH shifted from PPP genes *zwf* and *gnd* to the membrane transhydrogenase *pnt*, capable of direct transfer of electrons from NADH to NADP. Sauer *et al.* identified *pnt* as a major source of NADPH in *E. coli* (35-45% of total) [35]. Thus, *pnt* is capable of carrying significant flux *in vivo*. Taken together, these results suggested the model identified strains that promoted glycan precursor synthesis, primarily UDP-GlcNAc, by creating a combination of metabolite and redox imbalance.

**Figure 3:**
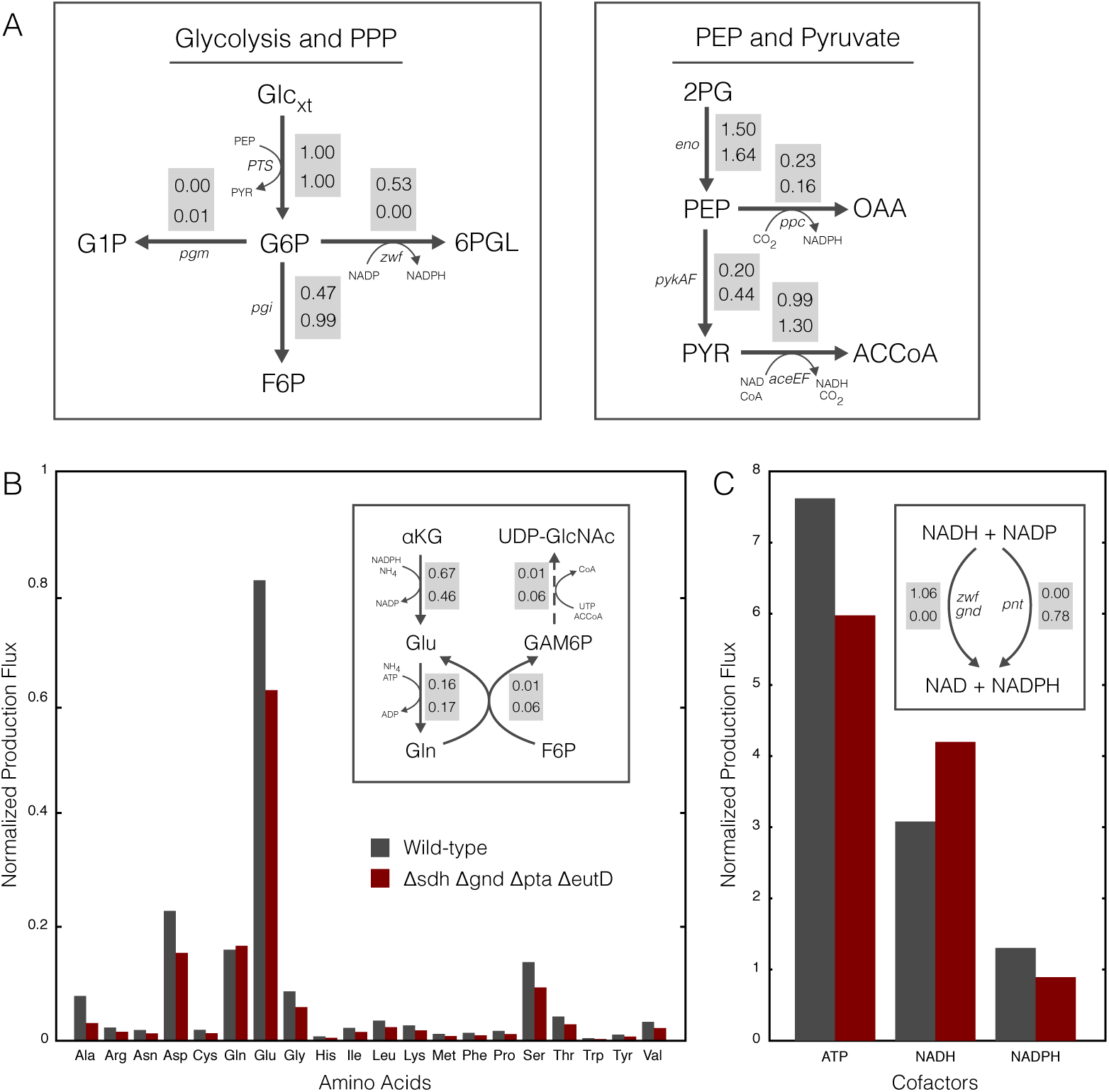
Comparison of fluxes between the wild-type case and glycan-producing strain of type EcGM3 as calculated by flux balance analysis. **(A)** Fluxes through key nodes of metabolism. Top fluxes correspond to the wild-type case, bottom fluxes are for strain EcGM3. Fluxes are normalized by the glucose uptake rate. **(B)** Total flux into each amino acid, normalized to glucose uptake rate. Inset shows fluxes associated with glutamate and glutamine synthesis along with the pathway to glycan precursor UDP-GlcNAc. The dotted arrow represents a lumped pathway of multiple enzymes leading to the glycan precursor. **(C)** Total flux into selected cofactors, normalized to glucose uptake rate. Inset shows the primary modes of NADPH production in each strain. Abbreviations: Pentose phosphate pathway, PPP; Extracellular glucose, Glc*_xt_*; Glucose-6-phosphate, G6P; Fructose 6-phosphate, F6P; 6-phospho D-glucono-1,5-lactone, 6PGL; Glucose 1-phosphate, G1P; Glycerate 2-phosphate, 2PG; Phosphoenolpyruvate, PEP; Pyruvate, PYR; Oxaloacetate, OAA; Acetyl-CoA, ACCoA; 2-Oxoglutarate, *α*KG; Glucosamine 6-phosphate, GAMP6P; UDP-N-acetyl-D-glucosamine, UDP-GlcNAc.

### Experimental validation of N-glycan-producing knockout strains

Glycan production was measured in the mutant strains to validate the model predictions (Fig. 4). Gene knockout strains were constructed using the Keio collection of single gene knockouts *E. coli* BW25113 [36] as donor strains for P1*vir* phage transduction. Mutants were constructed containing single, double, and triple knockouts that appeared in growth-coupled strains identified by the constraint-based model. We also performed simulations of each single gene knockout to determine genes that prevented glycan synthesis; *galU*, a key enzyme in the synthesis of glycan precursor UDP-glucose, was the only non-lethal knockout that prevented glycan synthesis. Knock-out strains were transformed with a plasmid constitutively expressing the *C. jejuni pgl* locus. To quantify glycan production, we took advantage of crosstalk between the glycosylation pathway and native lipopolysaccharide (LPS) synthesis in *E. coli* [37]. Specifically, after the glycan is flipped into the periplasm, it can be transferred to lipid A-core by the WaaL O-antigen ligase and shuttled to the outer membrane by LPS pathway enzymes, where it is displayed on the cell surface [16]. Labeling of these surface-displayed *N*-glycans with fluorescently-tagged lectins can then be used to quantify the amount of glycan displayed on the cell surface as a measure of glycan production [20]. Here, we labeled *C. jejuni* glycans for detection by flow cytometry with fluorophore-conjugated soybean agglutinin (SBA), a lectin specific to terminal galactose and GalNAc residues. Prior to labeling, knockout strains were grown in glucose minimal media and harvested during the exponential growth phase, to most closely satisfy the pseudo-steady-state assumption of model predictions.

**Figure 4:**
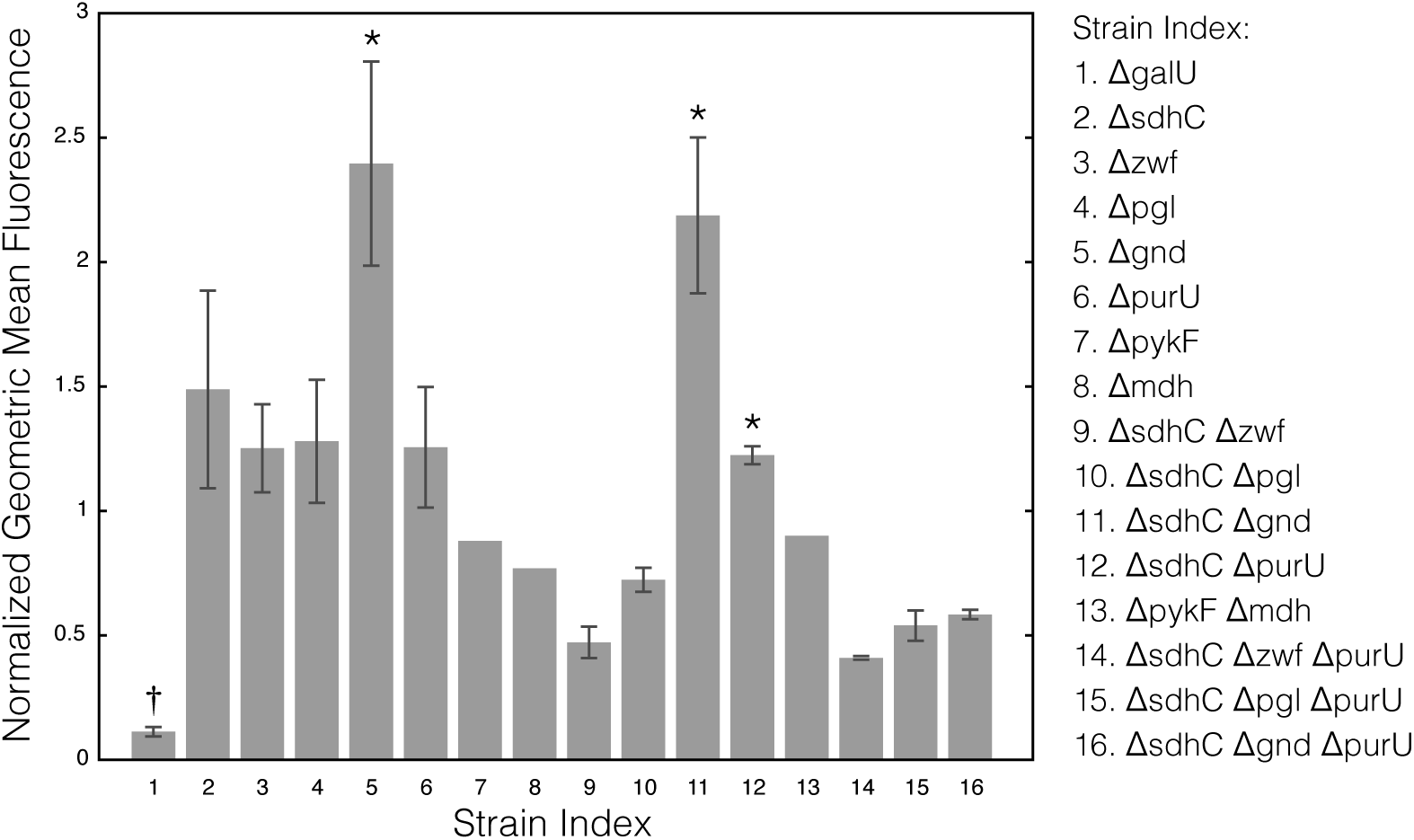
Geometric mean fluorescence, normalized to the wild-type value, from gene knockout strains appearing in growth-coupled strains identified by the constraint-based model. † indicates a strain predicted to eliminate glycan flux. Stars indicate statistically significant increases in fluorescences according to a t-test (*p <* 0.05). Error bars indicate the standard deviation of at least three replicates.

A common feature of the predicted mutant strains was the deletion of pentose phosphate pathway genes *zwf/pgl/gnd* in combination with Δ*sdh*. Analysis of the metabolic flux distribution in these mutants suggested the reducing state of the cell as well as the carbon flux was reprogrammed to support enhanced glycan biosynthesis. While hypothetical knockouts such as Δ*sdh* Δ*(zwf/pgl/gnd)* Δ*pta* Δ*eutD* were predicted to have higher glycan yield, in this study we experimentally evaluated only the simplest growth-coupled double knockout family, namely EcGM1. The EcGM1 family had the largest predicted growth rate, was more experimentally tractable than the triple and quad knockouts, and was an unambiguous test of the reducing power hypothesis without the complication of the additional deletions. Thus, while the EcGM2 and EcGM3 families could potentially give higher glycan flux, the EcGM1 family gave the clearest evaluation of the influence of the pentose phosphate pathway deletions. As predicted, single pentose phosphate knockouts Δ*zwf*, Δ*pgl*, and Δ*gnd* displayed greater fluorescence than wt cells, with Δ*gnd* being the most significant. However, when these deletions were combined with Δ*sdh* only the Δ*sdhC* Δ*gnd* combination led to increased glycan biosynthesis compared to wt cells. The single Δ*gnd* mu-tant increased glycan production by nearly 3-fold compared to the wt strain background, while the Δ*sdhC* Δ*gnd* combination led to a nearly 2.5-fold increase over the wt strain. Lastly, we tested the non-lethal deletions that were predicted to remove glycan biosynthesis; the Δ*galU* mutant showed no glycan production, thereby validating the model simulations. Taken to-gether, constraint-based simulations predicted pentose phosphate pathway deletions in combination Δ*sdh* (and potentially other genes) could improve glycan production by altering the redox state of the cell. We tested this hy-pothesis in the simplest possible experimental model, Δ*sdh* Δ*(zwf/pgl/gnd)*. Of the model predicted changes, only Δ*gnd* alone and Δ*sdhC* Δ*gnd* significantly increased glycan biosynthesis beyond the wt strain background. This suggested the model identified a potential axis for the improvement of glycan production, but results from the experimental system suggested this axis was likely more complicated as only the Δ*gnd* and Δ*sdhC* Δ*gnd* mutants gave a positive response.

## Discussion

In this study we adapted a genome-scale model of *E. coli* metabolism for the simulation of heterologous synthesis of *N*-glycans. We applied heuristic optimization in combination with flux balance analysis to identify genetic knockouts that coupled *C. jejuni* glycan synthesis to growth. Simulations identified growth-coupled strains for minimal media growth on glucose as the sole source of carbon and energy. Flux analysis of these strains revealed two modes of flux redistribution that promoted glycan synthesis. For growth on glucose, simulations showed that maintaining high glycolytic flux and producing excess glutamine for the amination of glycan precursor sugars led to a growth-coupled phenotype. Simulations also identified the PPP as a primary target, suggesting the manipulation of the NADH / NADPH ratio influenced glycan synthesis. We validated model predictions by measuring cell surface-displayed *N*-glycans in *E. coli* mutants. In all growth conditions, the Δ*gnd* mutant outperformed the wt strain in glycan synthesis. Over-all, our model-guided strategy showed promise toward rationally designing a microbial glycosylation platform.

We used simulated annealing and flux balance analysis to search for metabolic and regulatory gene knockouts that produced a growth coupled phenotype. Several constraint-based methods have been developed previously to identify gene knockouts that coupled production to growth e.g., [30, 38, 39]. Most of these methods rely on an OptKnock-like approach, whereby a bilevel mixed integer optimization problem is solved to identify the optimal set of gene knockouts. This class of method guarantees identification of the global optimum; however, it suffers from a few limitations. First, search time for OptKnock-like algorithms scales exponentially with system size and number of gene knockouts, making them unable to handle very large metabolic networks. Second, only linear engineering objectives (e.g., target production flux) can be searched over. In contrast, heuristic optimization is an effective approach for searching large networks while simultaneously considering non-linear objective functions. Though identification of the global optimum is not guaranteed with these methods, desirable sub-optimal solutions can be found quickly [38, 40]. Also, heuristic optimization can search efficiently for gene knockouts rather than reaction knockouts. This is an important distinction because the mapping of genes to reactions is not necessarily one to one. Thus, experimentally, many reactions may be difficult to knock out because they may be catalyzed by the products of many genes. Here, we used simulated annealing in combination with flux balance analysis to maximize the shadow price of growth with respect to glycan flux using a genome scale metabolic reconstruction. The approach identified PPP knock-outs that altered the NADH / NADPH balance, and increased glycolytic flux leading to enhanced glycan production. Surprisingly, these knockouts were not in the same section of the metabolism compared with previous literature studies. However, this may be expected, as we searched for growth coupled solutions and did not simply increase glycan formation. These solutions, while more difficult to obtain, offer a significant future advantage; namely, optimization of glycan production could be improved by selecting for increased growth through serial passage.

Many aspects of glycoprotein production in *E. coli* are amenable to investigation and engineering by metabolic modeling. This study focused on increasing the availability of glycan precursor metabolites through model-guided metabolic network manipulations. Other approaches in bacteria have focused on optimizing expression of glycosylation pathway enzymes and identification of metabolic reaction targets through proteomic and genome engineering [23, 26, 24]. Despite these efforts, improving glycosylation efficiency in *E. coli* remains a significant challenge. To address this challenge, a more comprehensive mathematical description of the cell, one that couples metabolism with gene expression and metabolic demand, may be required to precisely model glycosylation in *E. coli*. Our approach does not explicitly consider the metabolic burden associated with heterologous expression of glycosylation pathway enzymes nor the expression of the acceptor glycoprotein. Also, flux balance analysis lacks a description of enzyme kinetics and metabolite concentrations. Predicting phenotypic changes to genetic perturbations is a primary challenge in model-guided metabolic engineering [41]. It has been shown that single knockouts in the central metabolism of *E. coli* do little to change the relative flux distribution in the organism [42]. *E. coli* robustly controls metabolic flux using allosteric, transcriptional and post-transcriptional regulatory, and post-translational modification systems [43, 44]. Thus, glycoprotein production in *E. coli* is a unique challenge in that it requires optimization of two opposing cellular processes. Recombinant protein production of a desired glycoprotein along with glycosylation pathway enzymes requires energy from catabolic processes. On the other hand, glycan precursor synthesis requires conservation of available sugars and anabolic processes. The addition of regulatory systems and an explicit description of gene expression to a stoichiometric model may be an effective strategy for optimizing these opposing processes. Other strategies that may be helpful for optimization of this system include the enhancement of glycan precursor pathways, such as hexosamine synthesis, as well as the removal of competing pathways.

## Materials and Methods

### Flux balance analysis and heuristic optimization

Reactions encoding *C. jejuni* glycan formation (Table 1) were added to the genome-scale metabolic model of *E. coli* iAF1260 [27]. The combined model was then used to determine growth coupled gene knockouts that improved glycan production flux. Metabolic fluxes were estimated using flux balance analysis. Flux balance analysis requires two assumptions. First, the cell was assumed to operate at a pseudo-steady-state, where the rate of production of every intracellular metabolite was equal to its consumption. Second, the cell has evolved to operate optimally to achieve a cellular objective. Though many objectives have been proposed, we use the most common, namely growth rate (i.e., biomass formation) maximization [45]. The determination of a flux distribution satisfying these assumptions was formulated as a linear optimization problem:

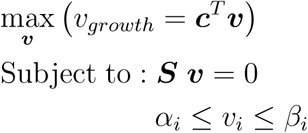

where ***v*** is the steady-state flux vector and *α*_*i*_ and *β*_*i*_ are the lower and up-per limits for the individual flux values, respectively. The quantity, *v*_*growth*_, denotes the specific growth rate where ***c*** is a vector containing the stoichio-metric contribution of each metabolic species to biomass. The stoichiometric matrix ***S*** encodes all biochemical reaction connectivity considered in the model. Each row of ***S*** describes a metabolite, while each column describes a particular reaction. The (*i, j*) element of ***S***, denoted by *σ*_*ij*_, describes how species *i* participates in reaction *j*. If *σ*_*ij*_ *>* 0, species *i* is produced by re-action *j*. Conversely, if *σ*_*ij*_ *<* 0, then species *i* is consumed by reaction *j*. Lastly, if *σ*_*ij*_ = 0, then species *i* is not involved in reaction *j*. The maximum substrate and oxygen uptake rates were set at 10 mmol/gDW/hr. Boundary conditions were set to allow for the unrestricted formation of acetate. All genes found to be essential for growth on Luria-Bertani (LB) medium were excluded from the search [36].

We used the FastPros algorithm developed by Ohno *et al.*, in combination with a shadow price objective, to estimate genetic knockouts [46]. Simulated annealing identified growth-coupled genetic knockouts with improved glycan production [47]. Prior to optimization, we removed all genes associated with dead end reactions, since knocking those out would have no effect on the network. Also, we removed duplicate genes, i.e., those that produced identical effects when knocked out. Finally, we removed genes whose knockout resulted in zero growth. We searched over both metabolic and regulatory genes; metabolic and transcriptional regulatory genes were represented by a binary array where 1 indicated the gene was expressed, and 0 zero indicated it was removed from the network (or transcriptionally repressed). A random initial gene knockout array was generated. We allowed for a maximum of 20 knockouts during the search. New knockout arrays were generated through crossover and mutation operators that randomly introduced new knockouts [38]. At each iteration, the fitness (shadow price) of an individual was computed using flux balance analysis. When an individual with a higher fitness was encountered (greater shadow price), that individual was accepted. However, when an individual with a lower fitness was encountered, we accepted this individual with a probability given by a Boltzmann factor:

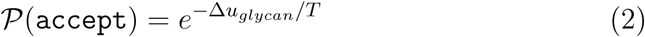

where Δ*u*_*glycan*_ denotes the change in shadow price between the current and previous solution, and the temperature *T* denotes the computational annealing temperature which decreased with the search iteration. The annealing temperature *T* decreased exponentially such that *T*_*k+1*_ = *αT_k_*, where *k* denotes the iteration index and *α* denotes the cooling rate defined as [40]:

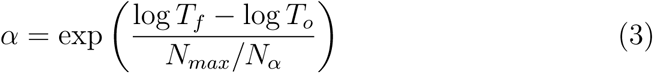

The term *N*_*max*_ denotes the maximum allowable number of objective function evaluations (*N*_*max*_ = 10, 000), and *N*_*α*_ denotes the number of objective function evaluations performed at each distinct temperature value (*N*_*α*_ = 1). The initial temperature *T*_*o*_was defined as 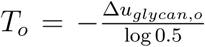, while the final temperature *T*_*f*_was given by 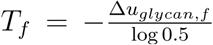. Lastly, Δ*u*_*glycan,o*_ denotes the difference in shadow price corresponding to an acceptance probability of worse solutions of 50% at the beginning of the search, and Δ*u*_*glycan,f*_ is the shadow price difference giving a 50% probability of accepting a worse solution by the end of the search. These values were approximated using the typical shadow price values of random knockout arrays: Δ*u*_*glycan,o*_ = 0.005, Δ*u*_*glycan,f*_ = 0.0005.

Though we sought to maximize glycan flux, we also wanted to identify experimentally viable strains. Thus, during an optimization search, we set a lower bound on the biomass reaction flux equal to 10% of the wild-type simulated growth rate. Strains that could not meet this constraint were ignored. The knockout search was terminated once a positive shadow price was found. After the optimization, we processed growth-coupled knockout strains by iteratively knocking in each knockout gene to estimate knockouts that did not affect the phenotype. In this way we identified the smallest number of gene knockouts that produced enhanced glycan flux at optimal growth. Each optimization run required approximately six hours on a single CPU Apple workstation (Apple, Cupertino, CA, USA; OS X v10.10). All model and optimization code is available in the MATLAB (The Mathworks, Natick MA) programming language, and free to download under an MIT software license from Varnerlab.org [29].

### Bacterial strains and media

For surface-labeled glycan fluorescence measurements, we used the *E. coli* strain BW25113 as our wild-type case [36]. BW25113 was used as the parent strain to construct all gene knockout strains. Plasmid pCP20 was used to excise KmR cassette [48]. Minimal media consisted of 33.9 g/L Na_2_HPO_4_, 15.0 g/L KH_2_PO_4_, 5.0 g/L NH_4_Cl, and 2.5 g/L NaCl. Media was supplemented with 0.4% glucose. Growth medium was supplemented by appropriate antibiotic at: 100 *μ*g/mL ampicillin (Amp), 25 *μ*g/mL chloramphenicol, and 50 *μ*g/mL kanamycin (Kan). Growth was monitored by measuring optical density at 600 nm (OD_600_).

### Flow cytometry

BW25113-based knockout strains were transformed with plasmid pACY-Cpgl, constitutively expressed by the *C. jejuni pgl* locus. Cultures were inoculated from frozen stock in LB and grew for 3-6 hours. Cells were subcultured 1:100 in minimal media overnight and then transferred to fresh minimal media to an OD_600_ of 0.1. 300 *μ*L cells were harvested during exponential growth phase (OD_600_ ≈ 0.6). Cells were washed with PBS then incubated in the dark for 15 minutes at 37*^°^*C. Cells were resuspended in 5 *μ*g/mL SBA-Alexa Fluor 488 (Invitrogen) and 500 *μ*L PBS and analyzed using a FACSCalibur (Becton Dickinson). Geometric mean fluorescence was determined from 100,000 events.

## Author contributions

J.V and M.P.D directed the study. J.W constructed the mathematical model and conducted the computational studies. T.M, C.G and J.W created the mutant strains and conducted the experimental studies. The manuscript was prepared and edited for publication by J.V, M.P.D, T.M and J.W. All authors reviewed the manuscript.

---

**Figure S1:**
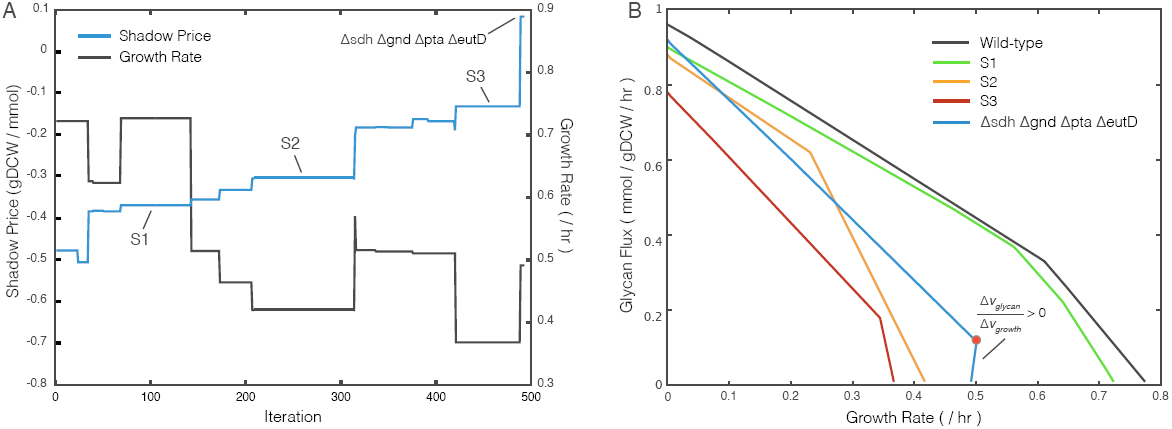
Representative simulated annealing simulation results identifying a growth-coupled strain of type EcGM2. **(A)** Shadow price and FBA-maximized growth rate versus iteration number for the identification of strain ΔsdhΔgndΔptaΔeutD. **(B)** Production envelopes of the strains (S1-S3) at the iteration points indicated in A. The simulation was terminated once a positive shadow price is found, visualized by the slope of the production envelope. The red dot indicates the optimal operating point of maximal growth rate for the growth-coupled strain.

